# Nanobody Detection of Standard Fluorescent Proteins Enables Multi-Target DNA-PAINT with High Resolution and Minimal Displacement Errors

**DOI:** 10.1101/500298

**Authors:** Shama Sograte-Idrissi, Nazar Oleksiievets, Sebastian Isbaner, Mariana Eggert-Martinez, Jörg Enderlein, Roman Tsukanov, Felipe Opazo

## Abstract

DNA-PAINT is a rapidly developing fluorescence super-resolution technique which allows for reaching spatial resolutions below 10 nm. It also enables the imaging of multiple targets in the same sample. However, using DNA-PAINT to observe cellular structures at such resolution remains challenging. Antibodies, which are commonly used for this purpose, lead to a displacement between the target protein and the reporting fluorophore of 20-25 nm, thus limiting the resolving power. Here, we used nanobodies to minimize this linkage error to ~4 nm. We demonstrate multiplexed imaging by using 3 nanobodies, each able to bind to a different family of fluorescent proteins. We couple the nanobodies with single DNA strands via a straight forward and stoichiometric chemical conjugation. Additionally, we built a versatile computer-controlled microfluidic setup to enable multiplexed DNA-PAINT in an efficient manner. As a proof of principle, we labeled and imaged proteins on mitochondria, the Golgi apparatus, and chromatin. We obtained super-resolved images of the 3 targets with 20 nm resolution, and within only 35 minutes acquisition time.

## Introduction

Super-resolution light microscopy is developing rapidly, and a growing number of cell biologists are embracing this technology to study proteins of interest (POI) at the nanoscale. Single molecule localization techniques like PALM[1], (d)STORM[2], [3], and others[4] achieve resolutions that allows for distinguishing molecules that are separated by only a few nanometers. Among these localization techniques, **DNA P**oint **A**ccumulation for **I**maging in **N**anoscale **T**opography (DNA-PAINT)[5] has demonstrated to achieve a resolution below 5 nm on DNA origami structures[6], [7] and offers the possibility to detect multiple POIs within the same sample[8]. A special feature of DNA-PAINT is that it is not limited by photobleaching of the fluorophore, due to the constant replenishment of fluorophores from solution. In fact, a target site carries one or more single stranded DNA oligonucleotides (commonly referred to as the docking strand or handle) instead of a single fluorophore, while a second single stranded DNA molecule with a complementary sequence to the docking strand bears a fluorophore (referred to as the imager strand). In a DNA-PAINT experiment, the imager strands continuously bind to the docking strands and unbind due to thermal fluctuations. The continuous transient binding of the imager strands results in sparse “blinking-like” fluorescence detection events. Similar to PALM or STORM, these events are then precisely localized to reconstruct a super-resolved image. The localization precision depends on the number of photons collected in a single event, whereas the total number of events recorded affects the quality of the final superresolved image. Importantly, DNA-PAINT benefits from the orthogonality of DNA hybridization (with different sequences). DNA docking strands with different nucleotide sequences can be associated with different targets, thus making it easy to obtain multi-target super-resolution images using a single fluorophore. Thereby, chromatic aberrations are avoided, resulting in a comparable resolution for all the POIs under investigation[9]. For such multiplexed imaging (known as Exchange PAINT[8],[9]), sequential introduction of different imager strands is required.

However, this methodology imposes several challenges to cell biologists who want to optimally image POIs with DNA-PAINT. Usually, primary antibodies that bind to a POI are labeled with secondary antibodies which carry the docking strand[10]. But such an approach introduces a spatial displacement of up to 25 nm between the target site and the fluorophore[11]–[13] which seriously limits the resolving power of all single molecule localization super-resolution techniques which use conventional antibody-based immunofluorescence labeling. The first attempt to minimize this “linkage-error”[14] was to use primary antibodies that are directly coupled to docking strands[15]. Typically, this has been performed by using an undirected coupling chemistry via maleimide-peG2-succinimidyl ester or via DBCO-sulfo-NHS-ester cross linkers[10]. These non-targeted coupling methods can interfere with the binding ability of the primary antibody to the POI by reacting at the paratope of the antibody. Additionally, they result in a mixture of antibodies containing a broad distribution of the number of docking strands (even including antibodies with none), which results in an inhomogeneous labeling density of the POIs and makes single molecule detection non-quantitative.

To further tackle the “linkage error” of the reporter fluorophores, several small monovalent affinity probes are continuously emerging[16]. For instance, small DNA or RNA molecules known as aptamers[17]–[19] or single-domain antibodies (sdAb, or nanobodies)[20] have recently gained popularity in the field of super-resolution imaging[21]–[23]. Nanobodies are obtained from a special type of immunoglobulins known as heavy chain antibodies (hcAb) and which are found in camelids. The recombinant production of the variable domain of these hcAbs result in a functional nanobody with only 2-3 nm size[24]. Recently, a significant improvement in spatial resolution, as compared to conventional antibody immunofluorescence, was demonstrated by using nanobodies for labeling[12], [25]. In addition to their small size, high specificity, and monovalent binding affinities, which make them an ideal tool for microscopy, the recombinant nature of nanobodies endows them with a great flexibility and allows introducing all types of modifications in a precise manner. This last feature permits to rationally design and control the number and location of desired functional elements on them (e.g. the number and locations of fluorophores or docking strands)[22].

Unfortunately, only few nanobodies able to recognize endogenous mammalian proteins are currently available. However, several new nanobodies against different fluorescent protein families like GFPs (from *Aequorea Victoria),* RFPs (from *Dicosoma* sp.) or mTagBFPs (from *Entacmaea quadricolor)* are now easily accessible. This opens the opportunity to obtain super-resolution images with a minimal linkage-error on a wide range of biological samples. Fluorescent proteins like EGFP[26], mCherry[27] and mTagBFP[28] are widely used in the life-sciences, fused to POIs within simple cell lines, large yeast libraries[29], and countless other genetically modified organisms (e.g. *Arabidopsis thaliana*[30], *Caenorhabditis* elegans[31], *Drosophila melanogaster,*[32] or mice[33]).

Here, we used a custom-built multi-channel Total Internal Reflection Fluorescence (TIRF) microscope and three nanobodies targeting mTagBFP, EGFP and mCherry to perform Exchange PAINT experiments on three different targets inside the same cell. For efficient buffer solution exchange, a versatile custom-built microfluidics system was developed and implemented. Exchange PAINT was performed by sequential introduction of three different imager strands and washing in between. Recorded single-molecule localization detection events were subsequently analyzed for reconstructing super-resolved images for each of the three targets. We achieved a resolution of 20 nm with a localization precision of 14 nm within 35 minutes of acquisition time (per target). We envision that nanobody-based DNA-PAINT will provide an efficient solution for the protein-DNA linkage problem and will help to exploit the full power of DNA-PAINT for cellular imaging, considering the broad availability of many fluorescent proteins.

## Materials and Methods

### Nanobody coupling to docking oligo

The unconjugated nanobodies FluoTag^®^-Q anti-GFP, FluoTag^®^-Q anti-RFP, and the FluoTag^®^-Q anti-TagBFP (NanoTag Biotechnologies GmbH, Cat. No: N0301, N0401, and N0501, respectively) carry one ectopic cysteine at the N-terminus then allowing for chemical couplings via a thiol reactive compound. The DNA docking strands (Biomers GmbH, Ulm, Germany) were functionalized with an azide group at 5′-end and, in some cases, Atto488 fluorophore at 3′-end. The coupling of the docking strands to the nanobodies were performed following procedure from Schlichthärle and colleagues[34], with minor modifications. In brief, 15 to 20 nmol of nanobodies in phosphate buffer saline (PBS, 127 mM NaCl, 10 mM Na_2_HPO_4_, 2.7 mM KCl, 0.2 mM KH_2_PO_4_, pH7.4) were incubated with a final concentration of 5 mM TCEP (Sigma-Aldrich) for 2 h at 4°C to reduce the ectopic cysteine. Afterwards, the excess of TCEP was removed by exchanging the buffer to PBS pH 6.5 using spin Amicon filters with a MWCO of 10 kDa (Merck/EMD Millipore, Cat. No. UFC501096). The reduced TCEP-free nanobodies were immediately mixed with 50 molar excess of DBCO-maleimide crosslinker (Sigma-Aldrich, Cat. No. 760668) and incubated overnight at 4°C with mild stirring. The excess of DBCO crosslinker in the buffer was exchanged to PBS pH 7.4 using Amicon Filters (MWCO 10 kDa) as described previously. Functionalized docking strands were added (10 molar excess) to the crosslinker-coupled nanobody, and incubated at room temperature for ~2 hours with slow head-to-tail shaking. The excess of docking strands was then removed from the conjugated nanobodies using size exclusion chromatography (Superdex 75 Increase 10/300 column, Cat. No: 29148721) and ÄKTA pure 25 system (GE life science). The correct fractions of labeled nanobodies were then identified by the SDS-PAGE followed by SYBR GOLD staining (Thermo Fisher, Cat No: S11494). The docking strands sequences used for the assay were taken from Agasti et al[8]. FluoTag^®^-Q anti-GFP was coupled to P1* sequence (5′-CTAGATGTAT-Atto488-3′), FluoTag^®^-Q anti-RFP was coupled to P2* (5′-TATGTAGATC-3′), and the FluoTag^®^-Q anti-TagBFP was coupled to P3* (5′-GTAATGAAGA-3′). Imager strands were labeled with Atto655 fluorophore at 3` end.

### Immunostaining

COS-7 cells were cultured in DMEM medium with 4 mM l-glutamine and 10% fetal calf serum (Thermo Fisher Scientific), supplemented with 60 U/ml of penicillin and 0.06 mg/ml streptomycin (Sigma-Aldrich) at 37°C and 5% CO2. Prior immunostaining and imaging, ca. 20,000 cells/well, were plated in 8-well chamber (155411PK, ThermoFisher Scientific). In the next day, the cells were transfected using 2.5% lipofectamine 2000^®^ and 300 ng of plasmid in Optimem medium (Thermo Fisher Scientific). After incubation of ca. 16 h, the cells were fixed using 4% paraformaldehyde (PFA) for 30 minutes at room temperature. The remaining aldehydes were quenched with 0.1 M glycine in PBS for 30 minutes. Afterwards, cells were permeabilized and blocked using 3% bovine serum albumin (BSA) and 0.1% Triton X-100 in PBS for 30 minutes at room temperature. Buffer solution containing nanobodies coupled to the docking strand (50 nM) was used to stain the cells. For this purpose, we proceeded with incubation of 1 h at room temperature, with slow orbital shaking. Finally, the cells were rinsed with PBS and then post-fixed with 4% PFA for 30 minutes at the room temperature. As described previously, remaining aldehydes were quenched with 0.1 M glycine in PBS. Cells were stored in PBS buffer at 4°C.

### Exchange PAINT Experiment

The imager strands P1 5′-CTAGATGTAT-3′-Atto655, P2 5′-TATGTAGATC-3′-Atto655, and P3 5′-GTAATGAAGA-3′-Atto655 (Eurofins Genomics) were aliquoted in TE buffer (Tris 10 mM, EDTA 1 mM, pH 8.0) at a concentration of 100 μM and stored at -20°C. Prior to the experiment, the strands were diluted to the final concentration of 2 nM in PBS buffer, containing 500 mM NaCl. A chamber with 8 wells (155411PK, Thermo Fisher Scientific) was fixed on the microscope stage with clips. A PDMS layer was used as a chamber cover and supported the inlet tubes and a tube for suction. The slide was held on the microscope stage for 0.5 h before the acquisition to equilibrate to the room temperature and achieve mechanical stability. Injection of fluids and its removal was done using our custom-built microfluidic setup, designed and constructed particularly for Exchange PAINT experiment. First, the well was rinsed twice with 500 μL PBS buffer (pH 8.0, NaCl 500 mM). Then, suitable cells for imaging were selected based on the presence of signal from the expressed fluorescent proteins: mTagBFP, mCherry, and EGFP. The cells were located by exposing them to the following laser excitation wavelengths and detecting the fluorescence in the corresponding emission channel: mTagBFP - 405 nm laser, EGFP - 488 nm laser, and mCherry - 561 nm laser. A HILO-illumination scheme was used to excite the cells. Laser power was adjusted according to the sample brightness (respectively 0.5 mW, 1 mW and 2 mW at the output of optical fiber). Each selection movie of the fluorescent proteins included between 200-250 frames (Figure 4, A1-A3). Afterwards, we proceeded with Exchange PAINT on the selected cell. All the solutions were injected into the cell by applying air pressure in the corresponding pressurized tube. First, imager strand P1 (2 nM) in PBS buffer (500 μL) was injected into the well and incubated for 10 minutes before the acquisition. Typical DNA-PAINT movie included 21,000 frames (corresponds to 35 minutes). The following acquisition settings for emCCD camera were used: exposure time 100 ms, pre-amplifier gain 3.0, EM gain 10. The laser 638 nm was set to 10-15 mW (corresponds to an excitation illumination power density of 0.4-0.6 kW/cm^2^). After PAINT movie acquisition, an extensive wash of the well was performed (4-6 times volume exchange, in total about 3 mL buffer within 5 minutes), in order to remove the imager solution from the well completely. Suction was performed by the micro peristaltic pump (Makeblock) After the extensive wash, the next imager solution was introduced. We proceeded with the same solution exchange procedure also for the imagers P2 and P3 (see comprehensive chart in Figure 3B). All the experiments were carried out at a constant temperature of 22±1 °C, which was crucial for the mechanical stability of the sample (remaining mechanical drifts were corrected for during the analysis).

### DNA-PAINT movies analysis

Raw DNA-PAINT movies were analyzed using the Picasso software package[10]. In the end, drift-corrected super-resolution images were reconstructed and the average localization accuracy was estimated. For further analysis of the achieved image resolution, the Fourier Ring Correlation (FRC)[35] technique was employed. First, localization events were detected using Picasso: Localize. For the specific binding-events recognition, the signal box size length was set to 7 pixels and the minimum net gradient was limited to the range of 1,700 to 3,500 (depends on the protein expression level in a particular cell). Then, the localized bright spots were fitted with the LQ Gaussian method to obtain precise fluorophore coordinates. The total number of localization events varied from 150,000 to 2,500,000 for the whole movie. The output file with the localization coordinates was then loaded into Picasso: Render. Using the Undrift RCC feature (segmentation 500 frames), movies were corrected for mechanical drift. The final reconstructed super-resolved images were exported in PNG format. Finally, all three reconstructed images of different organelles were merged together for each imaged cell using ImageJ[36], see Figure 4 C1,C2, D1-D3. The average localization precision (NeNa[37]) was estimated for each reconstructed super-resolved image. For image resolution quantification, Fourier Ring Correlation (FRC)[35] was applied using the FIRE ImageJ plugin [38], for detailed numbers see Supplementary Information, Table S1. Further image resolution analysis was performed by creating a resolution map using SQUIRREL[39] (super-resolution quantitative image rating and reporting of error locations), see Figure S4 in the Supplementary Information.

## Results

### Optimization of cells transfection and nanobody staining for Exchange DNA-PAINT imaging

First, we optimized the transfection of mammalian cells (COS-7) with plasmids encoding for proteins present in different organelles fused to various fluorescent proteins. We used TOM70 fused to EGFP to reveal mitochondria, GalNacT was fused to mCherry to detect the Golgi apparatus and histone H2B was fused to mTagBFP to detect the cellular chromatin (nucleus). Additionally, we used currently available nanobodies, which bind strongly and specifically to the three fluorescent proteins mentioned above. Each type of nanobodies was labelled with a unique docking strand, enabling the acquisition of multiple targets using Exchange PAINT, in single cells (see scheme in Figure 1).

**Figure 1.**
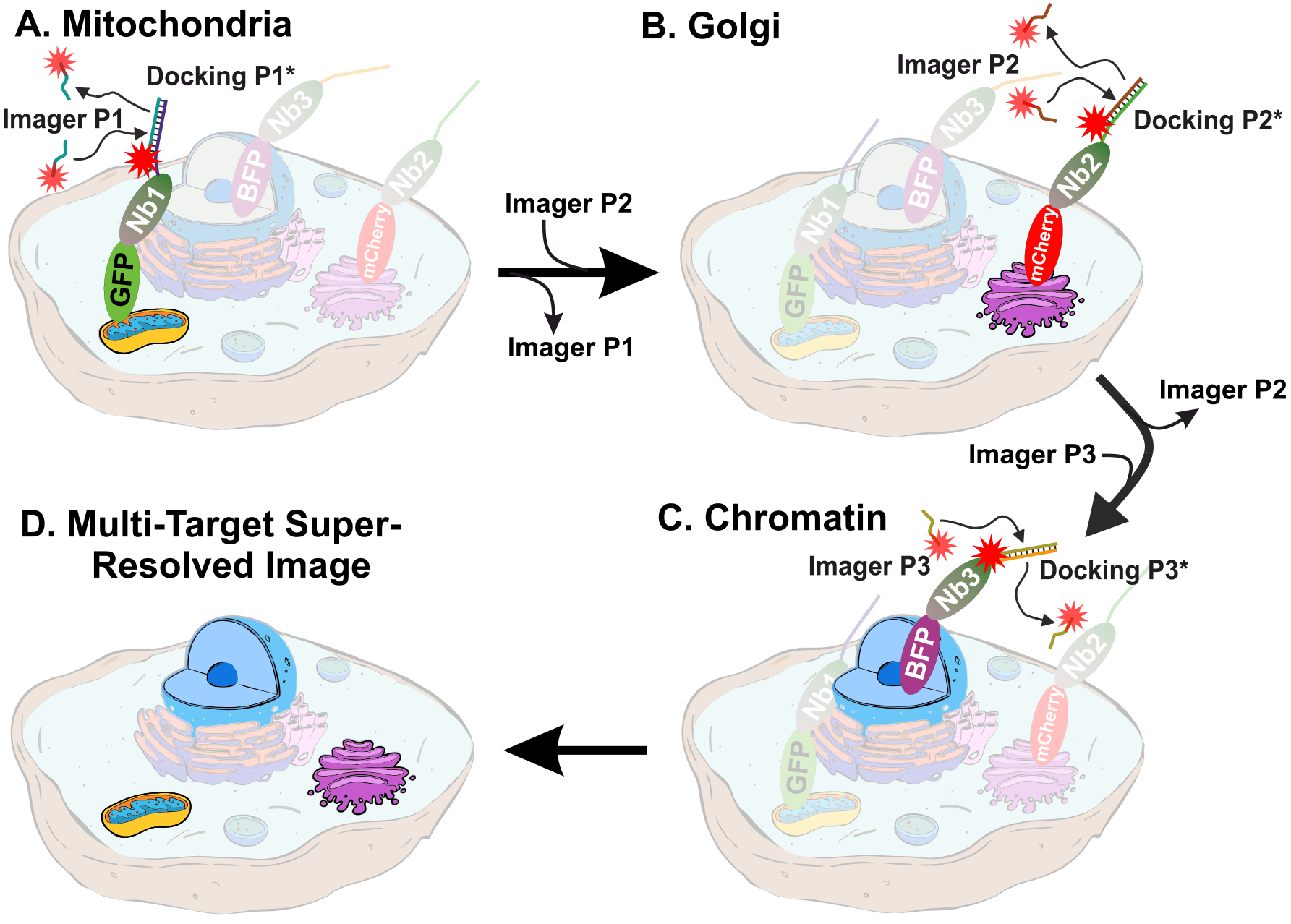
Schematic representation of multi-target Exchange PAINT in COS-7 cells. Sequential introduction of Imager strands with different sequences reveal multiple targets and result in multi-color super-resolution image. **(A)** DNA-PAINT imaging of mitochondria with Imager P1. **(B)** DNA-PAINT imaging of the Golgi apparatus with Imager P2. **(C)** DNA-PAINT imaging of nucleus/chromatin. **(D)** The resulting super-resolved image of a single cell with 3 colors overlaid. The cells were stained with (1) nanobody anti-GFP (Nb1) coupled to the DNA strand P1, (2) nanobody anti-mCherry (Nb2) coupled to the docking P2, and (3) nanobody anti-mTagBFP (Nb3) coupled to the docking strand P3.

All nanobodies had an extra ectopic cysteine at their C-terminus that allowed the conjugation of molecules via maleimide chemistry. We used a maleimide-DBCO as a cross-linker to attach the single stranded DNA oligo bearing an azide group on its 5′ end (Figure 2A, B). The coupling of the docking strand was thus performed in two sequential steps. First, the nanobody was incubated with a 50 molar excess of the maleimide-DBCO cross-linker, inducing a thiol-maleimide conjugation with the previously reduced single ectopic cysteine at the C-terminus of the nanobody[22]. After removing the excess of cross linker, the complex was incubated with a 10 molar excess of azide functionalized DNA oligo to induce a strain-promoted azide-alkyne cycloaddition (copper-free click chemistry[40]). The separation of the excess of DNA oligo from the mixture was performed using a size exclusion chromatography (SEC), resulting typically in two obviously separated elution fractions (Figure 2C). This is an essential step to avoid unspecific signal from the free DNA oligo. As a first routine quality control after SEC, different elution fractions were passed through a polyacrylamide gel electrophoresis (PAGE), stained with SYBR gold, to report for the presence of the oligonucleotides (Figure 2D). Only the fractions containing a clean band at the right molecular weight were used subsequently for the immunoassays of the transfected COS-7 cells. Due to the large excess of crosslinker and docking strands used for each coupling step (see Methods section), we are confident that a major proportion of the nanobodies were labelled with the docking oligo.

**Figure 2.**
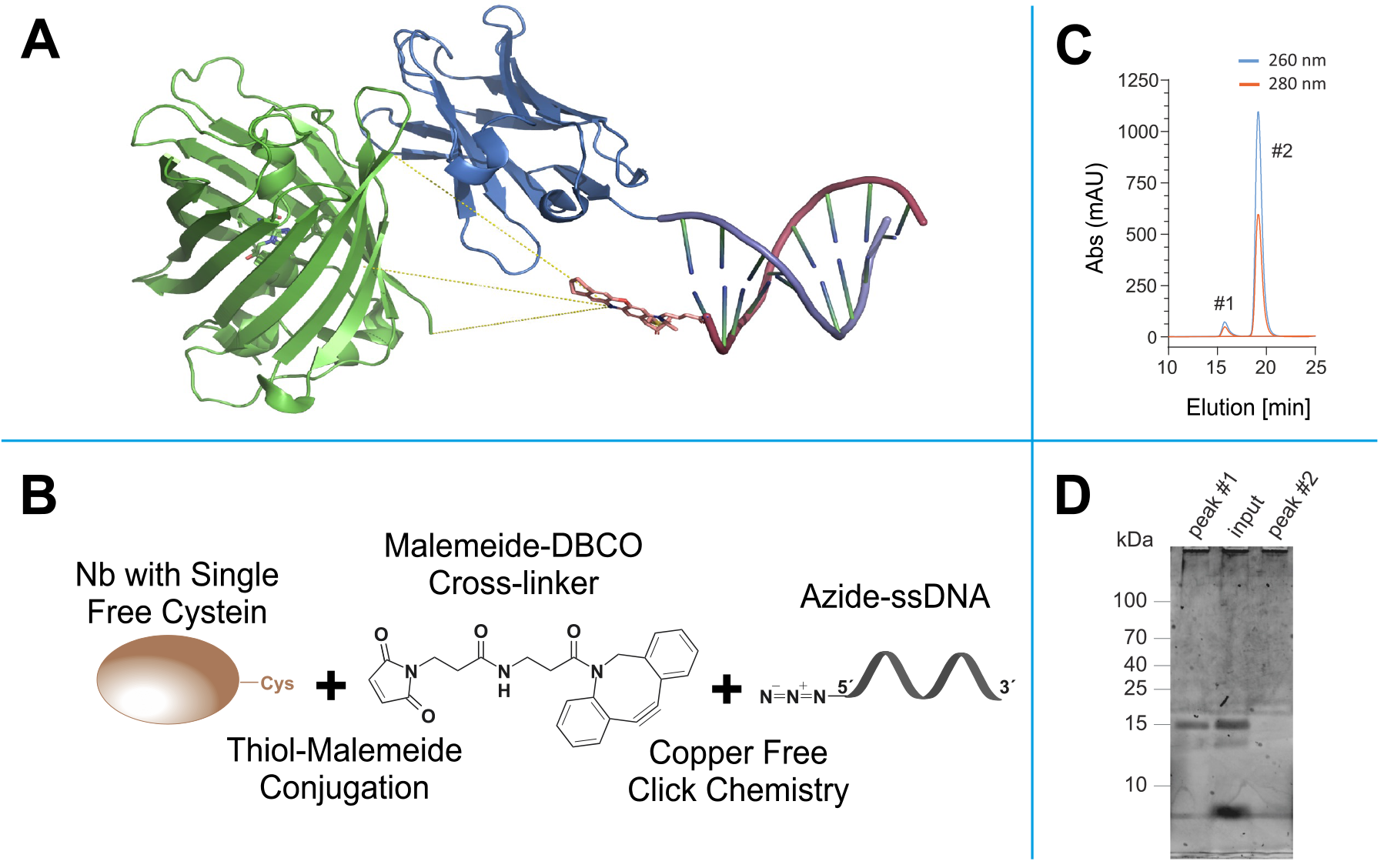
Click- and thiol-based strategy to conjugate nanobodies to a docking DNA strand for DNA-PAINT. **(A)** Anti-GFP nanobody (blue) bound to EGFP (green). The nanobody is modified with a docking stand with a complementary Atto655-labelled Imager strand attached (EGFP: nanobody complex extracted from (PDB: 3K1K), DNA strand and Atto655 were generated using ChemDraw (CambridgeSoft) and The PyMOL Molecular Graphics System (Schrödinger, LLC). Yellow lines represent 3 possible distances (3.1 nm, 3.3 nm and 3.4 nm) of the fluorophore to the POI. **(B)** Scheme representing the orthogonal coupling strategy of docking DNA strand to the nanobody. **(C)** Example of the size-exclusion chromatography (SEC) for the separation of DNA-coupled nanobody (#1) from the excess of docking strand (#2). **(D)** Example of the SDS-PAGE of fraction collected from the SEC run, post stained with SYBR gold, which reports DNA. Peak #1 collected from SEC shows a prominent band matching the expected molecular weight (~15 kDa). Peak #2 lacks the band at the nanobody molecular weight, suggesting that the peak contains the un-reacted excess of docking oligonucleotide.

### Microfluidic setup for Exchange DNA-PAINT experiment

Solution exchange inside the sample chamber was done using a custom-built microfluidics setup, designed and constructed in particular for the Exchange PAINT experiments. The setup allows operation of up to 24 independent inlet channels and is capable of fluid injection/removal with an adjustable flow speed in/out of the experimental chamber.

Magnetic solenoid valves (MH1, Festo) were used to turn on and off the air pressure in the channels, which were connected in turn to pressurized tubes. When air pressure is applied to such a tube, a liquid flowis created. Flow speed can be adjusted by changing air pressure using a pressure regulator (MS6, Festo). We used air pressure values in the range of 3-5 psi to generate a gentle flow of solutions. The pressurized air was purified with an air filter (PTA013, Thorlabs). Tygon tubing (VERNAAD04103, VWR) was used to guide the solutions from pressurized tubes to the experimental chamber. Suction was performed by a micro peristaltic pump DC12.0V (Makeblock). For washing, buffer solution for was loaded into a 15 mL test tube (Greiner Bio-One™ 188271, Fisher Scientific). The tube was equipped with a cap for pressurization (FLUIWELL 1C-15, Fluigent). The solutions of imager strands P1, P2 and P3 (concentration 2 nM, volume 1 mL) were loaded into 2.0 mL tubes (Micrewtube T341-6T, Simport), which were then mounted into a holder for four pressurized tubes (FLUIWELL-4C, Fluigent). Both magnetic solenoid valves and peristaltic pumps were operated by a custom-written LabView (National Instruments) routine, which included both manual and automatic operation modes. A custom-built electronic controller served as an interface between the magnetic valves and the computer.

### Super-resolution multiplexed DNA-PAINT images of COS-7 cells

We performed Exchange PAINT imaging of COS-7 cells stained with nanobodies, each functionalized with a single docking strand. For this purpose, aversatile custom-built optical setup was designed and constructed (see Figure S1). Initially, we checked that all cells to be imaged were triple transfected with the plasmids encoding for the TOM70, GalNacT and H2B fused to EGFP, mCherry and mTagBFP, respectively. The signal from each fluorescent protein was first imaged with a wide-field HILO illumination (see Figure 4, A1-A3). Afterwards, we sequentially introduced and removed imager strands P1, P2, and P3 as shown in Figure 3B. Each DNA PAINT movie was acquired during 35 minutes and then analyzed with the Picasso software[10] to obtain the super-resolved images (Figure 4, B1-B3). The experiment was designed to monitor three different proteins that are located in very distinct organelles, in order to simplify the evaluation of our Exchange DNA-PAINT images. Clearly, the reconstructed super-resolved DNA-PAINT images (one for each of the imagers P1, P2, and P3) showed the “patterns” expected for the respective organelle, thereby providing additional confirmation of our imaging strategy. For a more informative representation, and to further confirm the specificity of each imager strand, the three super-resolved images (from each P-imager movie) were merged together (Figure 4, C1). The result suggests that every “channel” remains clean, without significant unspecific binding events between the different docking-imager partners. The whole imaging cycle for the three target molecules, including imager strand injections, incubations, the removal of solutions and the acquisition of more than 60,000 frames took in total 2-3 h to be completed. The whole procedure worked robustly, and provided high quality superresolved DNA-PAINT images for nearly every imaged cell (selected at the initial wide-field HILO checkup).

**Figure 3.**
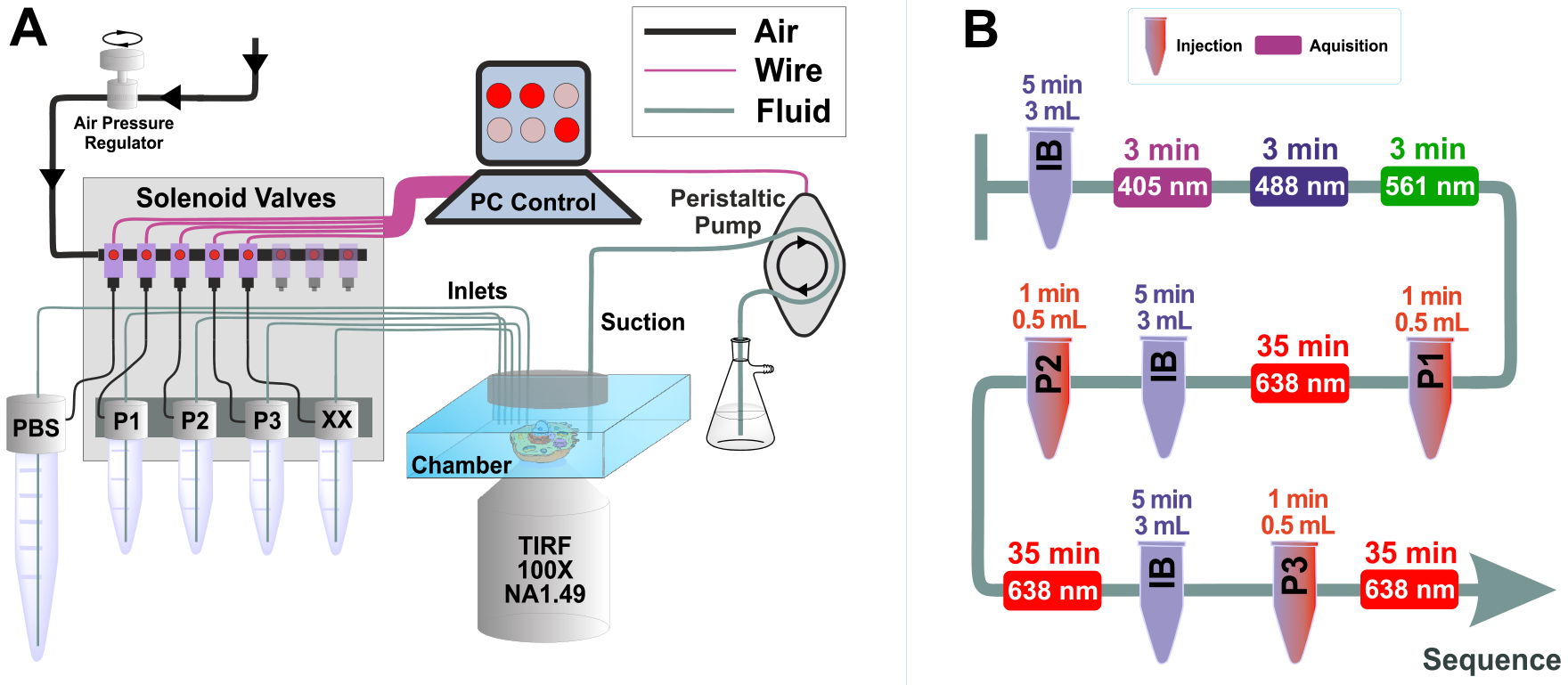
Scheme of custom-built microfluidics setup and Exchange PAINT experiment. (A) Microfluidics setup: the setup is controlled by a computer software, which includes both manual and automated operation modes. Maximum number of the input channels is 24 (only 5 channels are shown). The peristaltic pump used to remove the solutions from the chamber is also computer-controlled. (B) Typical sequence of actions for the Exchange PAINT experiment. The tube-shape sketches depict the injection of solutions (P1, P2 or P3) or the imaging buffer (IB) (solution volume and injection duration are indicated on top of the objects). Rectangles represent movie acquisition with certain laser excitation (laser wavelength and total acquisition time are indicated inside the rectangle and on top of the rectangle, respectively).

**Figure 4.**
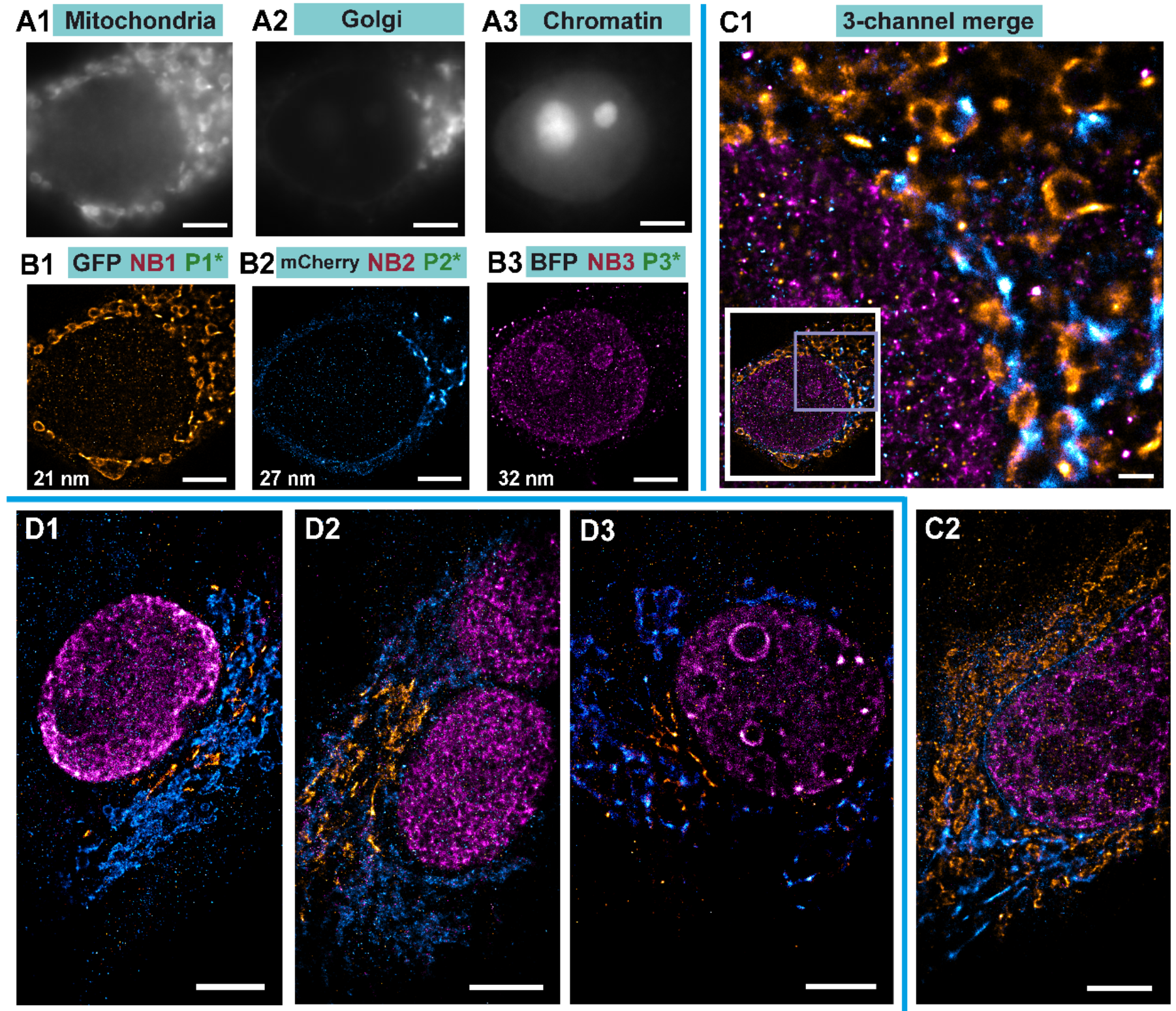
Exchange PAINT imaging. A1-A3 Diffraction-limited wide-field images of individual target fluorophores: Mitochondria with TOM70-EGFP (A1), Golgi with GalNacT-mCherry (A2), Chromatin with H2B-mTagBFP (A3). (B1-B3) Single-channel super-resolution DNA-PAINT images of respective organelles. The number represents FRC number for resolution for the respective image. (C1) Left bottom inset is the full view of the cell imaged in B with all 3-target Exchange PAINT images merged. Right, zoom of the boxed area I the inset. (C2) One more example of Exchange PAINT imaging with the same set of staining as in B and C. (D1-D3) Specificity controls were performed by swapping the fluorescent proteins and changing the Golgi marker. Now cells were expressing TOM70-mCherry and GM130-EGFP (a different Golgi marker). All scale bars correspond to 5 μm, except in C1 where the scale bar represents 1 μm.

In order to evaluate the image quality in a more quantitative manner, we performed a detailed analysis for the average localization accuracy and the actual resolution of the images. For the images presented in Figure 4, the average localization accuracy estimated by NeNa[37] was 19±2 nm (the lowest value was 14 nm, Table S1) and the average resolution, as estimated by Fourier Ring Correlation (FRC)[35], was 27±5 nm (the lowest value was 20 nm, Table S1). The full list of localization accuracies and FRC resolutions obtained for each organelle in each cell in Figure 4 can be found in Table S1 in the Supplementary Information.

Finally, we performed image quality analysis using FRC resolution maps using the SQUIRREL software[39]. The resolution values for different images varied between 26-34 nm. Moreover, we also compared the super-resolved images as obtained by Picasso and by RapidStorm[41]. Interestingly, both tools produce super-resolved images of similar quality (Figure S2).

To assess the effectiveness of our method, we performed several controls. First, the targeted protein GalNacT that we used to detect the Golgi, was changed to GM130 to ensure that the reconstructed organelle is labeled specifically regardless of the target protein used for reporting the Golgi apparatus (Figure 4, D1-D3). Additionally, we exchanged the fluorescent proteins we used for targeting the Golgi and the mitochondria by transfecting cells with plasmids encoding for TOM70 fused to mCherry and GM130 fused to EGFP. The same coupled nanobodies were used to reveal those targets. Nb1 (nanobody anti GFP) coupled to P1 docking strand, Nb2 (nanobody anti-mCherry) coupled to P2, and Nb3 (nanobody anti-mTagBFP) coupled to P3. Therefore, in this experiment, the imager P1 revealed the structure of the Golgi apparatus and the imager P2 revealed the mitochondria. Detailed comparison of NeNa and FRC for both cases can be found in Table S1. These control experiments confirmed the efficiency of our system, the specificity of the coupled nanobody, and the interchangeability of the targets. Finally, cells transfected with a single plasmid coding for TOM70-EGFP were immunostained with anti-GFP nanobodies bearing the P1-docking DNA and were imaged with imager P3 under the same conditions as before. We observed very few binding events (i.e. P3 imager binding P1 docking) without showing any recognizable pattern (Figure S3). This extra control suggests a high specificity of the imager to its docking strand. Additionally, unspecific binding (e.g. stickiness of the imager to the glass coverslip or cellular elements) were negligible (Figure S3).

## Discussion

Conventional antibodies (150 kDa and 12-15 nm in length) are often used for labelling cellular targets in DNA-PAINT imaging[5]. This approach has demonstrated to achieve an impressive spatial resolution[6], [7], to a level where the size of the primary and secondary antibody sandwich (with ~25 nm linkage-error) limits its imaging resolution. It has been demonstrated that small camelid single domain antibodies or nanobodies (15 kDa and ~3 nm in length) have the capacity to increase the accuracy of super-resolution microscopy for mapping POIs in a cellular context[15], [42]. Recently, nanobodies have been coupled to docking oligos to perform DNA-PAINT [25]. Unfortunately, only few nanobodies are currently available that work efficiently in immunoassay applications. Some of them have a strong affinity and high specificity towards specific fluorescent proteins. In this work, we exploited this property, which makes our method highly versatile since many bio-medical researchers typically have their favorite proteins already fused to fluorescent proteins[43]. Here, we showcase the use of three specific nanobodies against the EGFP family (this nanobody also binds to EYFP, Citrine, mVenus, Cerrulean, Emerald EGFP, and more GFP derivatives), mCherry and similar variants (it also binds to mOrange2, tdTomato, dsRed1 & 2, mScarlet-I, and other mRFP derivatives), and finally to mTagBFP (it also recognizes mTagRFP, mTagRFP657, mKate, and mKate2) for DNA-PAINT super-resolution microscopy. As a proof of principle, we used cells expressing three different fluorescent proteins in different organelles. The cells were immunostained with anti EGFP, mCherry, and mTagBFP specific nanobodies, each coupled to a unique and single DNA-docking strand for performing Exchange PAINT on them (Figure 4). We achieved an overall resolution of 20 nm, and an average localization precision of ~14 nm (in the best case, see Table S1), within only 35 minutes of acquisition time per target. We anticipate that, by further experimental and protocol optimizations, it will be possible to improve the resolution and acquisition time even further. The set of nanobodies presented in this work makes it already possible to investigate three proteins of interest within the same cell, all at diffraction-unlimited resolution, with the enhanced precision provided by the nanobody monovalency (no clustering of target protein) and small size (minimal linkage-error)[13], [16]. We hope that our study will motivate other scientists who have their POIs already fused to fluorescent proteins to benefit from this technique.

## Acknowledgments

The authors are grateful to Dr. Anna Chizhik for advising during the experiment, Dr. Oleksii Nevskyi and Arindam Ghosh for the fruitful discussions, Dr. Alexey Chizhik for advices in with graphics design and Dr. Ingo Gregor for the advices during the construction of the optical setup. We thank Dieter Hille and his team from the precision engineering workshop of the Third Institute of Physics, Georg August University. We also thank Markus Schönekeß and his team, in particular Simon Hoelscher, from the electronics workshop Third Institute of Physics, Georg August University. We thank Thomas Schlichthärle and his supervisor Prof. Dr. Ralf Jungmann for sharing their initial protocol and guidance for conjugation of the nanobody with the docking strand. We thank Dr. Selda Kabatas for helping with the molecular models in Figure 2. Finally, we would like to thank Prof. Dr. Silvio Rizzoli for his support.

