## Supplementary Information for "Nanobody Detection of Standard Fluorescent Proteins Enables Multi-Target DNA-PAINT with High Resolution and Minimal Displacement Errors"

**Running title:** Multicolor DNA-PAINT using nanobodies

**Keywords:** Nanobodies, Super-resolution microscopy, multi-color imaging, TIRF, microfluidics, DNA-PAINT, molecular localization, sdAb, GFP, mCherry, mtagBFP, multiplexing

**Custom-built optical setup description**

Measurements were performed on an in-house-built optical setup. In brief, the excitation part included four lasers: 405 nm (CUBE 405-100C, Coherent), 488 nm (PhoxX+ 488-100, Omicron), 561 nm (MGL-FN-561-100, Changchun), and 638 nm (PhoxX+ 638-150, Omicron). Manual shutters were used to easily switch between excitation lasers. The lasers were combined into the same optical path using dichroic mirrors DM1 (BrightLine DiO2-R561, Semrock), DM2 (BrightLine FF495-Di03, Semrock), and DM3 (zt405 RDC, Chroma). Then, the laser beams were coupled into a single-mode fiber (P1-460B-FC-2, Thorlabs) with typical coupling efficiency of 40-50%. After exiting the fiber, the beam was collimated and expanded by a factor of 3.6X using telescopic lenses. In order to achieve wide-field illumination, lens L1 (AC508-200-A-ML, Thorlabs) focused collimated laser beam on the back focal plane of the high-NA objective (UAPON 100x oil, 1.49 NA, Olympus). In order to switch between Epi-, HILO-, or TIRF-illumination schemes, the translation stage TS (LNR50M, Thorlabs) was used to mechanically shift the corresponding optical elements. The translation XY stage (M-406, Newport) ensured smooth and stable sample movement during the scan.

A separate translation stage with a differential micrometer screw (DRV3, Thorlabs) was holding the objective and was used for focusing. The emitted fluorescence was separated from the excitation laser using the multi-band dichroic mirror DM4 (Di03 R405/488/532/635, Semrock). Lens L2 (AC254-200-A-ML, Thorlabs) was used as a tube lens. An adjustable slit (SP60, OWIS) was positioned in the image plane and was used to limit the field of view. The multi-band filter BP1 (ZET488/561/635m, Chroma) was used to filter out laser remains in the detection path. Lenses L3 (AC254-100-A, Thorlabs) and L4 (AC508-150-A-ML, Thorlabs) were used to transfer the image plane from the slit to the EMCCD camera (iXon Ultra 897, Andor), thereby providing rectangular space for wavelength-based splitting of the emission light into two or three emission channels according to the experimental requirements. For this purpose, the dichroic mirrors DM5 (Chroma 550 LPXR) and DM6 (FF648-Di01, Semrock) were positioned on magnetic bases MB (KB50/M, Thorlabs). For each channel, additional band-pass filters were used: BP2 (BrightLine FF 445/20, Semrock) for the blue channel, BP3 (BrightLine FF 536/40, Semrock) for the green channel, and BP4 (BrightLine HC 692/40, Semrock) for the red channel. The overall magnification factor of the optical setup was 166.6X, the pixel size was 103.5 nm x 103.5 nm and the full field of view was 53 μm X 53 μm. Focus stability was achieved by robust construction of the custom microscope body, tightly fixing the 8-well chamber (155411PK, ThermoFisher Scientific) to the sample holder and keeping the temperature in the room stable.

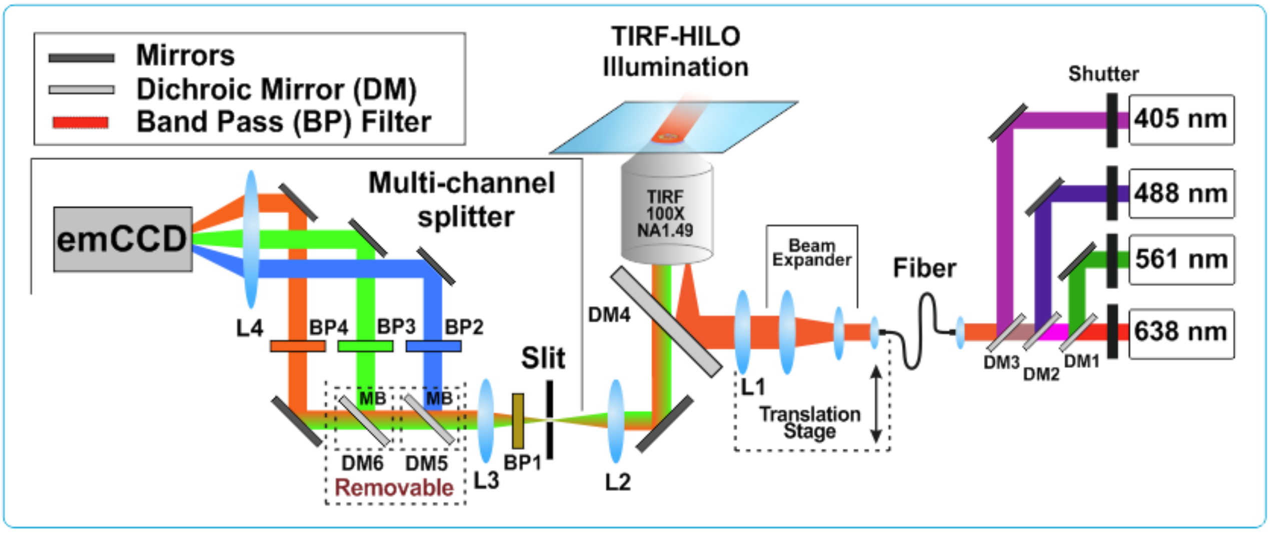

**Figure S1. Schematic drawing of custom wide-field TIRF optical setup**. The excitation is equipped with four lasers and allows excitation of fluorophores on broad spectral range. Multi-channel detection enables simultaneous imaging of fluorophores with different emission spectrum on the same CCD camera. Number of emission channels can be easily switched between one, two and three channels by removing the dichroic mirrors DM5 or/and DM6. The size of region of interest is controlled by a slit.

### Localization accuracy and resolution estimations

Detailed numbers as for each reconstructed image resolution are listed in the Table S1.

| Cell name | Organelle, protein, docking | NeNa, nm | FRC, nm |
|---|---|---|---|
| Fig 4, C1 | Mito GFP P1* | 20 | 21 |
| | Golgi RFP P2* | 18 | 27 |
| | Nucleus BFP P3* | 22 | 32 |
| Fig 4, C2 | Mito GFP P1* | 22 | 34 |
| | Golgi RFP P2* | 16 | 23 |
| | Nucleus BFP P3* | 23 | 34 |

| Cell name | Organelle, protein, docking | NeNa, nm | FRC, nm |
|---|---|---|---|
| Fig 4, D1 | Mito RFP P2* | 19 | 22 |
| | Golgi GFP P1* | 18 | 24 |
| | Nucleus BFP P3* | 23 | 38 |
| Fig 4, D2 | Mito RFP P2* | 20 | 28 |
| | Golgi GFP P1* | 19 | 20 |
| | Nucleus BFP P3* | 19 | 24 |
| Fig 4, D3 | Mito RFP P2* | 14 | 24 |
| | Golgi GFP P1* | 18 | 27 |
| | Nucleus BFP P3* | 17 | 26 |

**Table S1. Average localization precision (NeNa) and resolution estimation using FRC technique for each of the reconstructed images presented in Figure 4.** Cell names appear according to the order in Figure 4.

### DNA-PAINT analysis toolkits comparison: Picasso vs. Rapidstorm

Verification of Picasso analysis was done by analysing same image with RapidStorm.[1] Recorded DNA-PAINT movie was loaded into RapidStorm. Blinking event were identified by setting the intensity threshold to 60 % of a total brightness. Resolution (both X and Y direction) was set to 10 nm/pixel. The comparison between reconstructed images shown on Figure S2. The achieved resolution was estimating by exporting the localization file output and then running it into SQUIRREL ImageJ plugin[2]. Resulting numbers and comparison to Picasso output shown on Table S1. We saw good agreement Picasso and RapidStorm results.

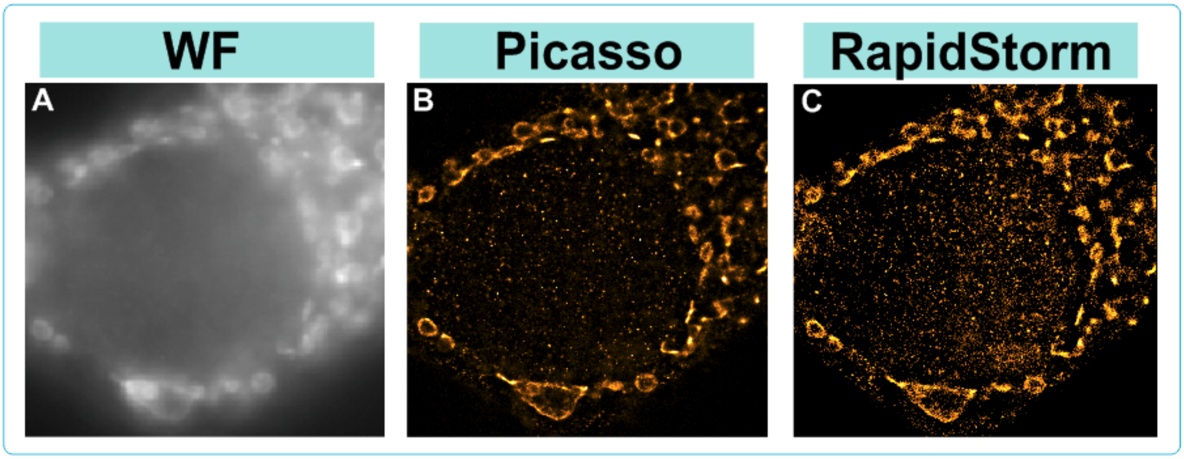

**Figure S2. Comparison between Picasso and RapidStorm software packages.** (**A**) Wide-field (WF) diffraction-limited image of GFP protein. (**B**) Reconstructed super-resolution image obtained with Picasso software. (**C**) Reconstructed super-resolution image obtained with Picasso software.

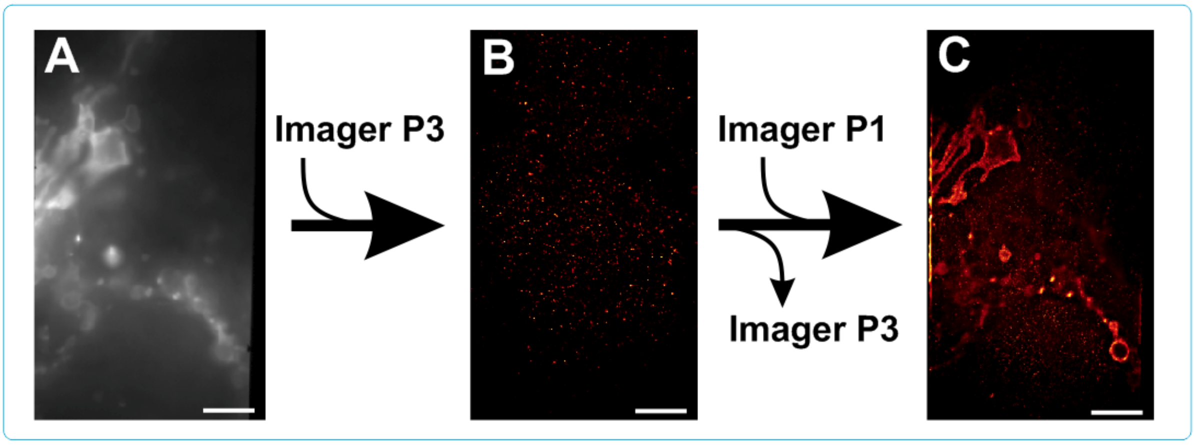

**Figure S3. Control experiment for the nanobody specificity and the stickiness of the imager strand inside fixed COS7 cell transfected with TOM70-GFP-P1***. (**A**) wide-field image of TOM70 GFP. (**B**) Super-resolution image from a DNA-PAINT movie taken in presence of P3 imager. Only few random localization events detected in presence of the imager P3 (**C**) Super-resolution image of the same region of interest, taken in presence of P1 imager (after washing/removing the imager P3).

### **DNA-PAINT Fourier Ring Correlation resolution maps**

For NanoJ-SQUIRREL analysis stack of two statistically independent super-resolution images of the same structure were reconstructed using Picasso and RapidStorm software. Then using ‘Calculate FRC Map’ feature (block per pixel value 25 and pixel size 10 nm) the FRC map was made and overlaid with respective super-resolution image. Average FRC resolution value was obtained by finding the mean value from the area with high localization density. For this purpose, obtained super-resolution image was cropped and whole procedure was repeated for this area. The average resolutions obtained are 26 nm for Picasso and 30 nm for RapidStorm.

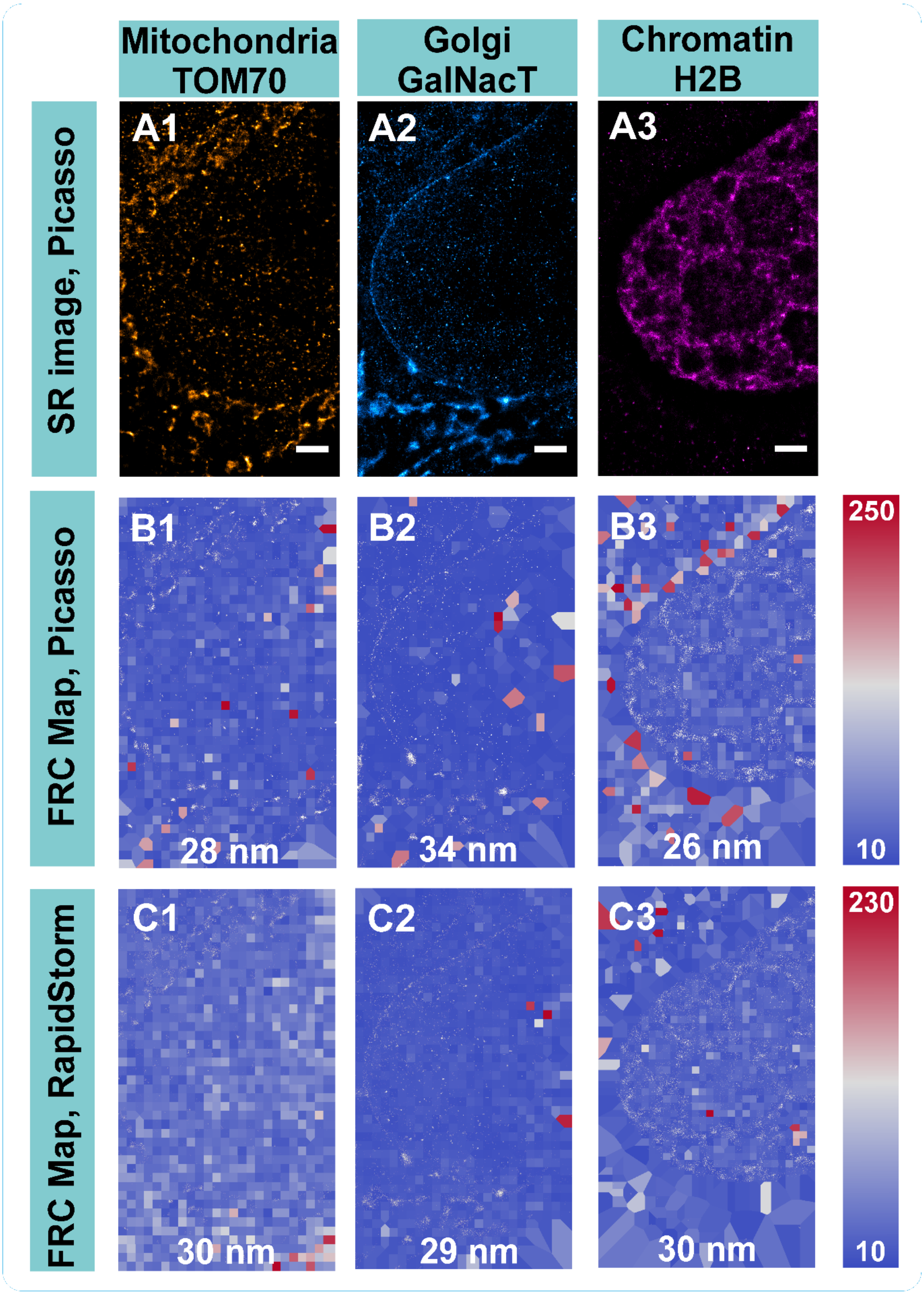

**Figure S4. Resolution estimation using Fourier Ring Correlation maps realized by SQUIRREL ImageJ plugin[2].** Comparison between Picasso and RapidStorm analysis tools. (A1-3) Super-resolution DNA-PAINT images of COS7 cell organelles reconstructed by Picasso. (B1-3) FRC map overlaid with the corresponding super-resolution images, as reconstructed by Picasso toolkit. (C1-3) FRC map overlaid with the corresponding super-resolution images, as reconstructed by RapidStorm toolkit. The numbers on the images represent the resolution averaged over the object region. The colors on B and C indicated resolution, according to scale shown on the right-hand side.
